# Competition and niche partitioning of floral resources between two native stingless bees (*Melipona mimetica* and *Scaptotrigona* sp., Apidae: Meliponini) in a seasonally dry tropical forest of Ecuador

**DOI:** 10.64898/2026.03.29.715153

**Authors:** Bruna Vieira, Francisco Lopes, Daniel Griffith, Elizabeth Gusman, Carlos Ivan Espinosa

## Abstract

Stingless bees are key pollinators in tropical ecosystems, yet their ecological dynamics remain poorly understood in highly seasonal environments such as the seasonally dry tropical forests of Ecuador. These ecosystems experience pronounced climatic seasonality, with sharp transitions between dry and wet periods that strongly affect floral resource availability. Understanding interspecific competition and niche partitioning in such systems is critical, particularly given the global decline of pollinators. We investigated resource use and niche dynamics in two native stingless bees, *Melipona mimetica* and *Scaptotrigona* sp., by quantifying pollen, nectar, and resin collection across seasons. Log-linear models were used to test the effects of species, season, and their interaction on resource use, while non-metric multidimensional scaling (NMDS) assessed niche overlap. Contrary to the expectation that niche overlap increases under resource scarcity, we found greater overlap during the wet season, when resources are more abundant. This suggests that both species converge on high-quality floral resources during peak availability, reflecting an adaptive response to strong environmental seasonality. Pollen use remained stable across seasons, consistent with generalist foraging behavior. In contrast, nectar collection increased significantly during the wet season, while resin exhibited a shared seasonal peak, likely associated with synchronized nest construction or maintenance. These findings reveal context-dependent competition dynamics and highlight the role of environmental seasonality in shaping pollinator interactions. Our study provides new insights into the ecology of threatened stingless bees and contributes to their conservation in tropical dry forest ecosystems.

## Introduction

Understanding the biological mechanisms that enable species coexistence has long been a central goal in community ecology. In environments characterized by a high diversity of ecologically similar species and limited resources, classical ecological theories (i.e., Gause’s principle and Hutchinson’s niche theory; Gause, 1934; Hutchinson, 1957) predict that competition leads to the exclusion of some species. However, many of these species coexist, raising the question of how this coexistence is achieved. Niche partitioning, often shaped by competitive interactions, provides a key mechanism whereby species exploit resources across spatial, temporal, and/or functional dimensions, reducing niche overlap and facilitating coexistence (Pickett et al., 2018; Petalas et al., 2021), often expressed as resource partitioning (Schoener, 1974).

For pollinating animals such as bees, this mechanism is particularly relevant. Most bees forage individually on patchily distributed floral resources in highly competitive environments (Dubois et al., 2021), where competition is driven by limited resource availability, high diversity, and frequent niche overlap (Roulston & Goodell, 2011; Page & Williams, 2023). Accordingly, niche partitioning is widely documented in bee communities (e.g., Casanelles-Abella et al., 2023; Ishii et al., 2008; Ye et al., 2024; Dubois et al., 2021), with coexisting species reducing competition through differences in floral preferences, morphological traits, foraging time, and spatial resource use.

Niche partitioning is particularly pronounced in seasonal environments, where marked climatic seasonality, which is characterized by strong contrasts between dry and rainy periods, drives pronounced fluctuations in resource availability and environmental conditions across time (Silveira et al., 2016). Such conditions are found in seasonally dry tropical forests (SDTF) in southern Ecuador, which are defined by two well-marked climatic seasons: a prolonged dry season lasting approximately eight months and a shorter rainy season of around four months (Espinosa et al., 2012). These ecosystems exhibit high levels of endemism in both plants and animals (Vázquez et al., 2005) and, consequently, most ecological processes are strongly seasonal (Aguirre et al., 2006), requiring species to adapt to pronounced temporal fluctuations in resource availability.

Among the diverse fauna inhabiting these forests are stingless bees of the tribe Meliponini, one of the most abundant and ecologically important groups of pollinators in tropical ecosystems (Maliani, 2006; Balvanera, 2012). Such is the peculiarity of these insects that the dynamics of said forest and other crops depend on the activities that these organisms’ practice (Rasmussen, 2004). According to Roubik (1989), the high diversity of Meliponini reflects their extensive participation in food webs and their importance as pollinators, particularly in comparison to abiotic pollination, i.e., wind and water.

As generalists, stingless bees obtain resources from a diverse group of plant species (Acereto, 2008), yet their foraging activity is often concentrated on a limited subset of plants, influenced by the availability and spatial distribution of floral resources (Cortopassi et al., 2006). The perennial and social nature of stingless bee colonies promotes continuous foraging, including the collection of nectar, pollen, and resins from a wide range of plant species (Vossler, 2012), contributing to their generally low degree of specialization (Flores et al., 2015). However, despite this generalist tendency, foragers frequently exhibit floral constancy and temporary specialization, concentrating their activity on a limited number of plant species at a given time. This pattern is driven by foraging efficiency, central-place constraints, and information exchange within colonies, such as recruitment through jostling behaviour, rather than by resource scarcity alone (Ramalho et al., 2007). Consequently, while temporary specialization may increase local competition when multiple colonies converge on highly rewarding resources, it can also promote niche partitioning across spatial and temporal scales, as different colonies or species exploit distinct subsets of floral resources, thereby reducing overall niche overlap.

Several studies suggest that analysis of fodder pollen loads as a useful tool to evaluate bees’ diets given their dependence on flowers (Sommeijer et al., 1983; Ramalho et al., 1990). Sommeijer et al. (1983) found that pollen variability is greater between *Melipona* species than among colonies of the same species, suggesting that, although resource use varies among colonies, interspecific differences are more pronounced, indicating stronger niche partitioning at the species level. Likewise, similarity in pollen diets may vary according to the availability of floral resources (Nagamitsu et al., 1999). Eltz et al. (2001) and Nagamitsu et al. (1997) affirm that foraging patterns tend to reflect the diversity of resources available within a given area, without marked differences, and that coexistence among species may be facilitated by interspecific differences in resource use.

Resource partitioning in stingless bees has been documented, yet information on these interactions remains limited in Ecuador, particularly in seasonally dry tropical forests. For this reason, in the present study we compare the type and composition of floral resources collected by two native species, *Melipona mimetica* ‘bermejo’ and *Scaptotrigona* sp.’catana’, to assess patterns of resource partitioning and potential competition across seasons. Both species are widely used for honey production, which is highly valued for its nutritional and medicinal properties, in southern Ecuador, especially in Loja Province. Taking advantage of the spatial arrangement of managed colonies and concurrent flowering dynamics during the sampling period, we evaluated whether resource use is structured by competition or niche partitioning between these two species. We hypothesized that temporal variation in resource availability would play a stronger role than interspecific differences in shaping resource use, potentially reducing the importance of niche partitioning in this seasonal ecosystem.

## Material and Methods

### Study site and data collection

The study was conducted in the La Manga sector of Garza Real parish, located in Zapotillo Canton, Loja Province, southwestern Ecuador (UTM coordinates: 17S 577174.86 E, 9532505.85 N; 17S 577624.3 E, 9532430.64 N). The region is dominated by an SDTF ecosystem, characterized by lowland dry scrub vegetation and ranging in altitude from 525 to 547 m.a.s.l. This ecosystem has been classified as critically endangered (Ferrer-Paris et al., 2018). The region climate is tropical semiarid with an average mean annual temperature varies 24°C and annual precipitation fluctuates between ca. 660 and 1300 mm (Espinosa et al., 2018). The climate is strongly seasonal, with a dry season from May to December and a rainy season from January to April. During the dry season, mean temperatures reach up to 26 °C and mean monthly precipitation is around 10 mm, whereas in the rainy season temperatures increase to about 28ºC and precipitation rises substantially to around 100 mm (Espinosa et al., 2018). The area is characterized by vegetation dominated dry seasonal forests with deciduous and semi-deciduous species, with pronounced leaf loss during the dry season. Flowering is strongly seasonal and typically associated with rainfall, leading to marked temporal variation in floral resource availability.

### Study species

The two focal species, *Melipona mimetica* and *Scaptotrigona* sp., were selected due to their ecological relevance and their commercial and cultural importance in Ecuador. Both species commonly coexist in the same habitats and exploit similar floral resources (Gaona et al., 2019), making them suitable models for investigating resource partitioning and potential competition. These species differ in key biological traits, including foraging range, morphology, and resource collection (e.g., pollen and resins), which may influence their patterns of resource use. *Melipona mimetica* is generally larger and more robust than *Scaptotrigona* sp., reflecting differences commonly observed between these genera. In addition, differences in nest structure have been reported, with *Melipona* species generally producing larger honey pots than *Scaptotrigona* (Vit et al., 2023).

### Sampling design and data collection

Data on floral resource use and pollen composition were collected in June, August, October, and December 2015, and in May 2016, covering a full annual cycle including both dry and rainy seasons. We sampled three colonies per species in each of two meliponaries (managed stingless bee breeding sites), separated by approximately 500 m. Within each meliponary, colonies were spaced approximately 3 m apart. To characterize the available floral resources, a circular transect of approximately 2 km was established around the meliponaries, corresponding to the average foraging range of the study species. Flowering plants were surveyed along this transect, and samples of floral buds were collected for botanical identification using reference collections. For each colony, 30 foraging individuals were sampled per sampling event. Before sampling, the entrance of each colony was temporarily obstructed using a net to increase the accumulation of returning foragers near the nest entrance. Foragers were captured using an entomological net and manual collection between 08:30 and 14:00 h. The resource carried by each individual was identified in situ as pollen, nectar, or plant resin. Pollen loads were removed from the corbiculae using cotton swabs, placed in labelled 2 ml Eppendorf tubes, and stored for subsequent analysis. Individuals carrying nectar or resin were recorded but not sampled.

### Pollen processing

Corbicular pollen was processed in the laboratory following the Wodehouse (1935) pollen preparation method. Additionally, the natural method described by Maurizio (1975) was used, in which a glycerogelatin solution was prepared using 7 g of gelatin, 50 ml of glycerin, and 1 g of phenol diluted in 42 ml of distilled water. For stained samples, fuchsin was added to the glycerogelatin solution. Pollen material from each individual was diluted in 95% ethanol on microscope slides. The cotton swab containing the pollen was gently scraped, and the material was divided into three replicates from the same individual bee. After ethanol evaporation, 2 μl of glycerol with fuchsin stain and 2 μl without stain were added to each slide. The acetolysis method was not applied; therefore, some morphological characteristics of the pollen may not have been fully resolved.

Pollen analysis was conducted using digital photographs obtained from prepared slides under a ZEISS-AxioStar Plus optical microscope with a 40× objective. The central area of each slide was systematically scanned, and five photographs were taken per section (i.e., with and without fuchsin staining). Pollen grains were identified following the nomenclature proposed by Erdtman (1952), using reference databases, identification guides, and comparison with a reference pollen collection (palynoteca) from dry forest vegetation associated with the study area. For resource quantification, the presence–absence of each pollen type was recorded for both stingless bee species, and the corresponding percentages were calculated. Similarly, the proportion of individuals carrying nectar and plant resins was determined. Data were analysed by sampling period to allow temporal comparisons of bee–plant interactions, and a binomial generalized linear model was used to assess differences in resource use between species.

### Statistical analysis

Resources are first analysed and organized as a multi-way contingency tables and secondly analysed using log-linear models. We want to know if the variation of resources is dependent on time, species or both. Our null hypothesis it that the predictor and response variables are not associated with each other, i.e. the two variables are independent of each other. We create 9 models. The model 8 of three-way interaction is compared with 9, which is the full model or Akaike. After this, variables are diminished, and the conditional independence is analysed by comparing models 7, 6 and 5 with 8, and finally the marginal independence is analysed by comparing models 4, 3 and 2 with model 1. In this analysis, the model that best explains the results is chosen, and therefore has the lowest AIC with the least number of variables. The models are compared using ANOVAs. All analysis was developed in R studio (RStudio team, 2019). To analyse the compartmentalization of the niches, according to the type of pollen that each species of bee *Melipona mimetica* and *Scaptotrigona sp*. they load in each sample, NMDS (non-metric multidimensional scaling) are performed in R (RStudio team, 2019) for each sample. We use the Jaccard distance, index that is based on presence absence, and the Morisita distance, index that considers abundances. To analyse the differences in the core and the total homogeneity of the niche of each species, we use the Adonis and betadisper function of the vegan package (Oksanen et al. 2019). Adonis compares similarity values of the centroids of the niches, while betadisper compares data about the homogeneity and size of the niches. The results of these are checked by ANOVA tests.

## Results

A total of 251 individuals of *M. mimetica* and 262 of *Scaptotrigona* sp. were recorded, of which 200 and 187 individuals, respectively, carried pollen, resin, or nectar loads. In total, 1,539 pollen samples were analysed. Across the six colonies, a total of 68 pollen types were recorded, corresponding to 31 families, 38 genera, 32 species, and three spore types (Table S1). Differences in pollen use were evident based on presence–absence data. Overall, both species exhibited similar patterns, with no significant differences between species. Model 4, selected as the most parsimonious model, indicated that resource use varied significantly over time, independent of species (Table 1). Figure 1 illustrates temporal variation in resource use by *M. mimetica* and *Scaptotrigona* sp.

**Table 1.** Candidate models assessing the effects of species, resource type, and sampling period. Model selection was based on Akaike’s Information Criterion (AIC), with Model 9 representing the full model. Model 4 was selected as the most parsimonious model.

| MODEL | G-squared | df | P value | AIC |
| --- | --- | --- | --- | --- |
| 1. frequency, species + resource + sampling | 65.253 | 22 | 0 | 65.253 |
| 2. frequency, species + resource + sampling + species: resource | 64.41 | 20 | 0 | 64.410 |
| 3. frequency, species + resource + sampling + species: sampling | 65.253 | 18 | 0 | 65.253 |
| 4. frequency, species + resource + sampling + resource: sampling | 12.162 | 14 | 0.593 | 12.162 |
| 5. frequency, species * resource + species * sampling | 64.41 | 16 | 0 | 64.410 |
| 6. frequency, species * resource + resource * sampling | 11.319 | 12 | 0.502 | 11.319 |
| 7. frequency, species * sampling + resource * sampling | 12.162 | 10 | 0.274 | 12.162 |
| 8. frequency, species * resource + species * sampling + resource * sampling | 11.303 | 8 | 0.185 | 11.303 |
| 9. frequency, species * resource * sampling | 0 | 0 | 0 | 0 |

**Figure 1.**
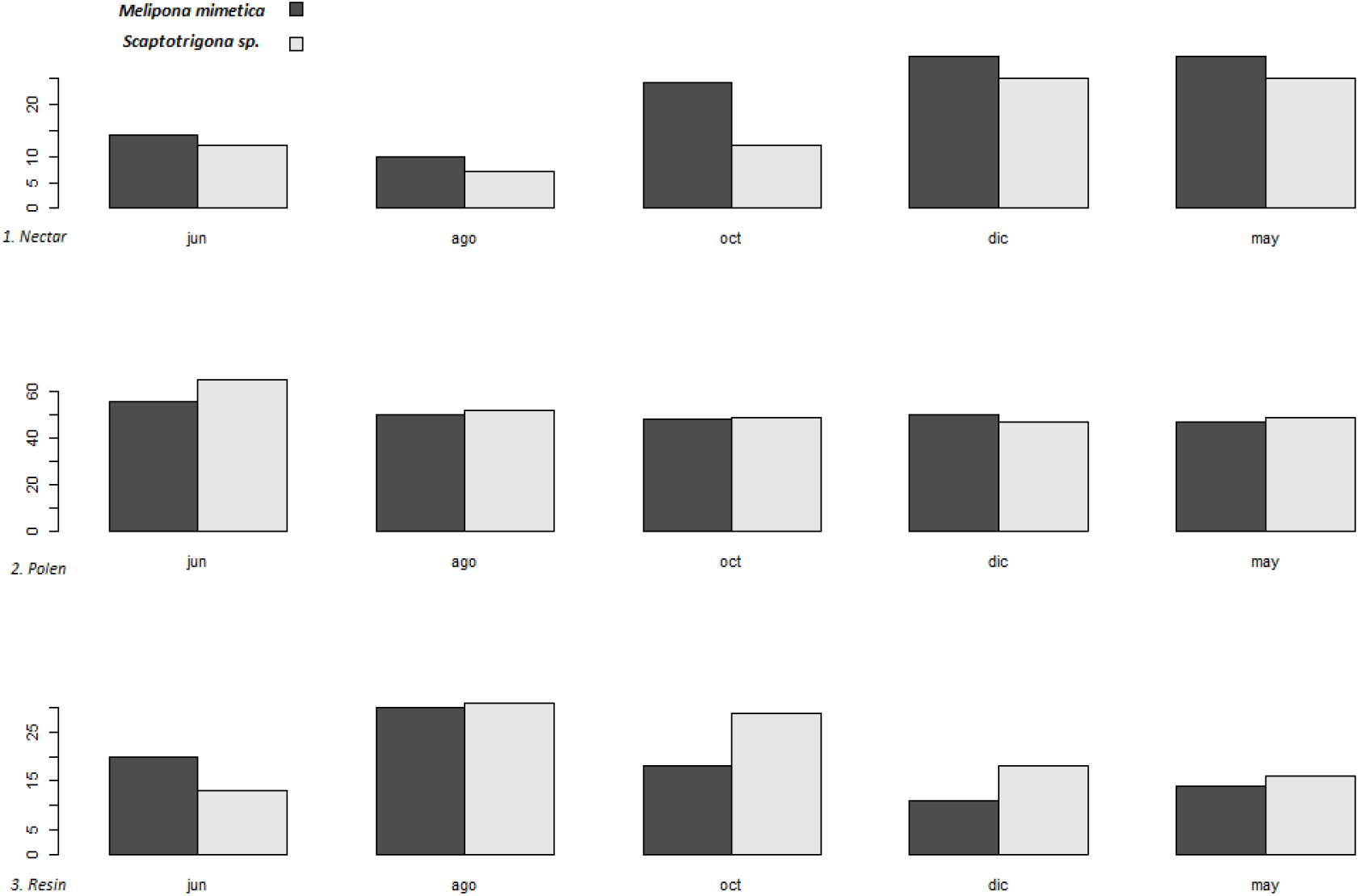
Use of resources (nectar, pollen and resin, from the top to bottom) by Melipona mimetica (color black) and Scaptotrigona sp. (color gray), along the five periods of sampling (June, August, October, December and May)

Resource use varied over time (Figure 2). Nectar showed a U-shaped pattern, with lower values in intermediate periods and higher values at the beginning and end of the sampling period. Pollen remained relatively stable, with a slight decline over time. In contrast, resin showed a unimodal pattern, peaking at intermediate sampling periods.

**Figure 2.**
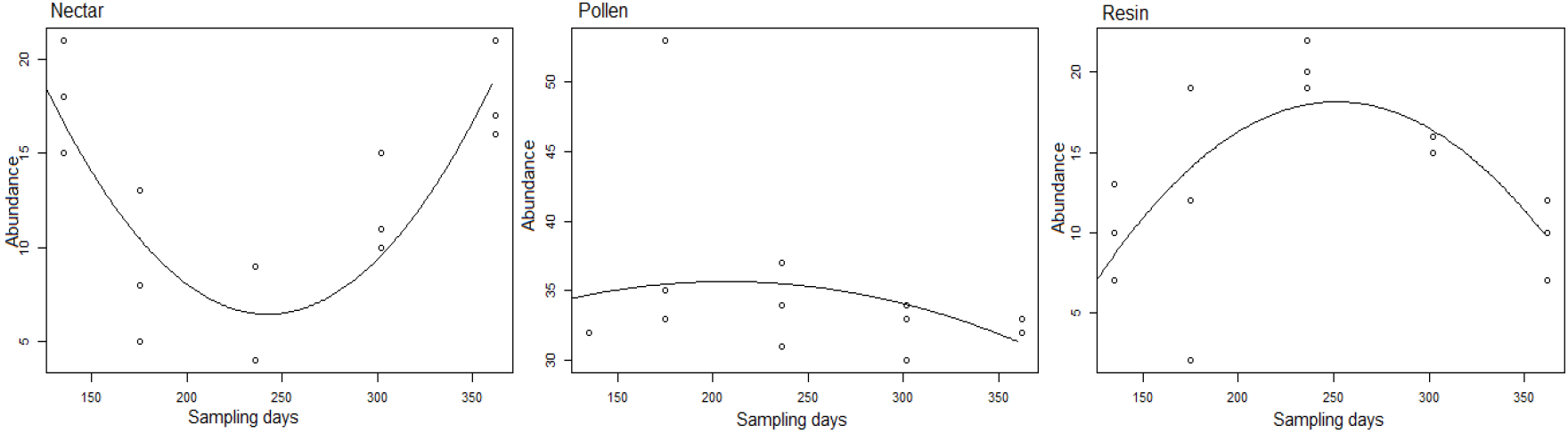
Abundance variation of resources (nectar, pollen, resin) throughout the year in which the sampling was carried out. As shown in the graphs, nectar and resin resources vary throughout the year, however pollen remains practically stable throughout the year

NMDS ordinations indicated temporal variation in pollen composition (Figure 3). The overall ordination showed a moderate fit (stress = 0.26), whereas period-specific analyses showed improved representation (stress = 0.13–0.23). A general overlap between *M. mimetica* and *Scaptotrigona* sp. was observed across sampling periods, although partial separation occurred in some periods (e.g., June and August), suggesting temporal shifts in resource use.

**Figure 3.**
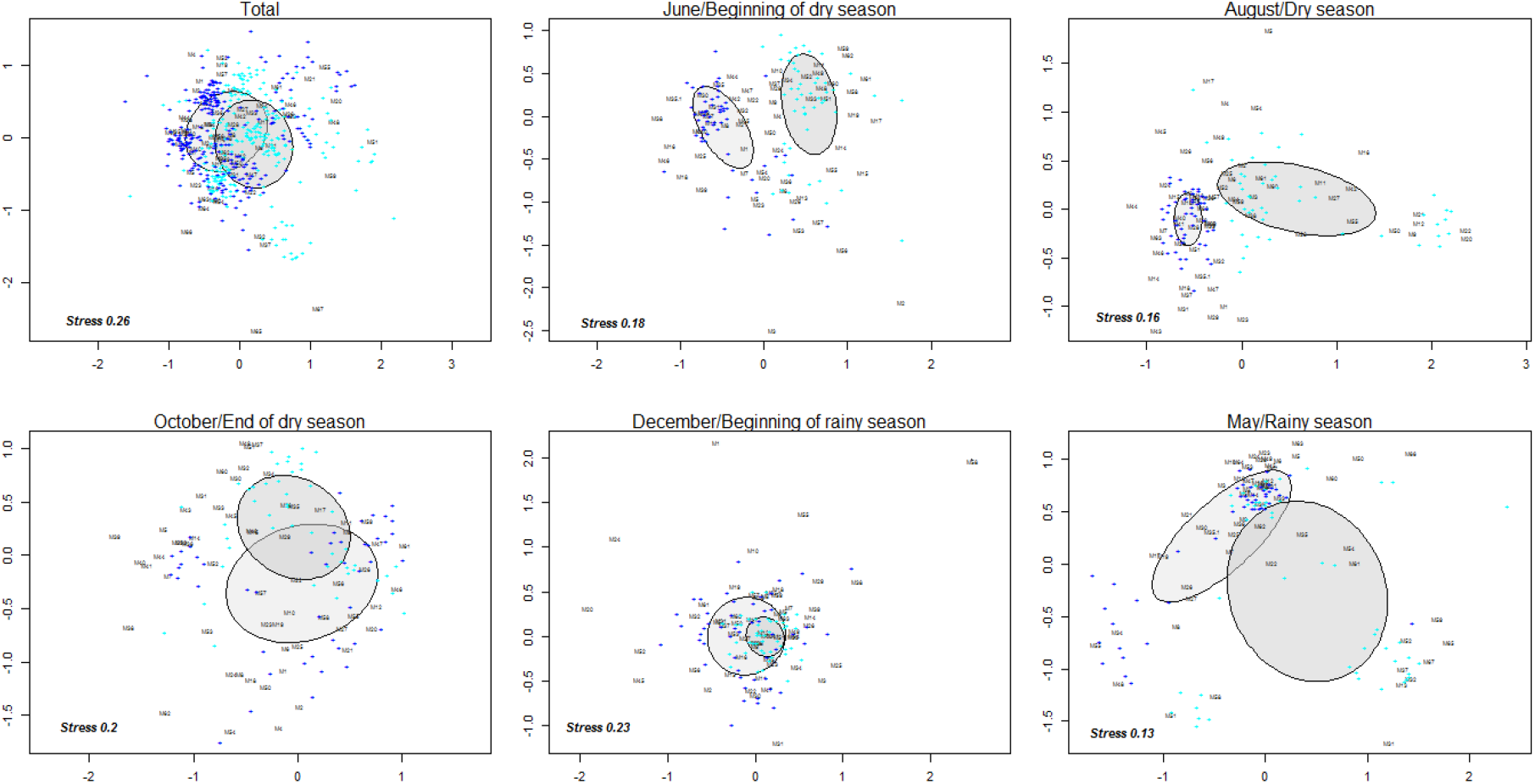
NMDS analysis using Jaccard distance for the niche partitioning of Melipona mimetica and Scaptotrigona sp. Its shows in the first plot the total of all samplings, and then the plots of individual samples of June (beginning of dry season), August (dry season), October (end of dry season), December (beginning of rainy season) and May (end of rainy season).

## Discussion

Results of the present study show that resource use by both stingless bees – *Melipona mimetica* and *Scaptotrigonas* sp. – was primarily driven by temporal variation rather than species-driven effects. Different resource types exhibited distinct seasonal patterns, with nectar and resin showing marked temporal variation, while pollen remained relatively stable. Despite these fluctuations, both species exhibited broadly similar patterns of resource use, with substantial niche overlap that varied across sampling periods, consistent with previous studies on stingless bees (Eltz et al., 2001; López-Roblero et al., 2023; Bernhardsson et al., 2024). While niche overlap is often associated with periods of resource scarcity in some systems (e.g., temperate ecosystems; Casanelles-Abella et al., 2023), our results revealed a different pattern: overlap increased during the rainy season, suggesting that greater floral availability promotes convergence in resource use.

These patterns can be understood within the broader ecological context of tropical dry forests, which exhibit an intermediate level of seasonality between highly seasonal temperate systems and aseasonal tropical rainforests. Such seasonality strongly influences floral resource availability and, consequently, the structure of species interactions. In highly seasonal systems, environmental constraints can restrict pollinator activity and promote temporal or functional niche differentiation (Slater et al., 2004; Mohra et al., 2004), whereas in aseasonal tropical forests, niche partitioning may occur along alternative axes, such as vertical stratification or floral traits (Nagamitsu et al., 1999). In contrast, tropical dry forests combine elements of both systems, leading to dynamic patterns of resource use and niche overlap across seasons.

In terms of resource use, nectar availability declined throughout the dry season, reached a mid-season minimum, and increased again with the onset of rainfall, consistent with patterns reported in tropical ecosystems where precipitation strongly regulates plant phenology and floral resource availability (van Schaik et al., 1993; Borchert et al., 2004). Such patterns are also captured by vegetation indices such as NDVI, which show marked seasonal variation in tropical dry systems, increasing from approximately 0.4–0.6 during the dry season to 0.6–0.8 in the rainy season, reflecting substantial increases in vegetation greenness and productivity (e.g., Viana & Célia, 2011). This temporal pattern was consistent across both species, with nectar use increasing similarly during the rainy season (Fig. 1).

Pollen use remained relatively stable across sampling periods, suggesting a relatively continuous availability of pollen resources throughout both dry and rainy seasons. The observed pattern likely reflects the diversity of pollen sources within the plant community, the generalist foraging behaviour of stingless bees (Acereto, 2008) and the asynchronous flowering phenologies characteristic of tropical dry forests (Frankie et al., 1974; Borchert et al., 2004). Such temporal complementarity ensures a relatively continuous supply of pollen, even during periods of low overall floral availability. Both species showed a similar peak in resin use during august (Fig. 1), suggesting that during this period the two share colony-level requirements. Moreover, the essential role of pollen as a primary protein source for colony development (Heinrich, 2004; Pang et al., 2022) may further contribute to its consistent use across seasons.

In comparison, resin use exhibited marked temporal variation, with a peak during intermediate sampling periods (Fig. 2). Unlike nectar and pollen, resin is not a nutritional resource but serves essential functions in nest construction and defense (Shanahan & Spivak, 2021), and its use is therefore not solely determined by environmental availability. Instead, resin foraging is strongly influenced by colony-level conditions, such as developmental stage and colony strength (Ferreira Junior et al., 2010), meaning that temporal peaks may reflect internal colony demands. The observed increase in resin use may correspond to periods of intensified nest construction or maintenance, as reported in previous studies. Consistently, both species showed a similar increase in resin use during this period (Fig. 1), suggesting that this pattern is not species-specific but likely driven by shared colony-level requirements.

Ordination analyses (NMDS) revealed that niche overlap between species varied across sampling periods, closely tracking seasonal changes in resource availability and use (Fig. 3). During the rainy season, greater overlap was observed, particularly in relation to nectar, suggesting convergence in resource use when floral resources are abundant. A possible explanation for this pattern lies in optimal foraging theory, which predicts that foragers preferentially exploit the most profitable resources when these are widely available, thereby reducing the energetic costs associated with resource search (MacArthur & Pianka, 1966; Pyke et al., 1977), and allowing them to concentrate on the most rewarding resources when available, promoting convergence in resource use. Under such conditions, multiple species may converge on the same high-reward floral resources. This pattern may be further reinforced in tropical systems during periods of high flowering intensity – such as the seasonally dry tropical forests of Ecuador -, when abundant resources attract a wide range of foragers (Frankie et al., 1974; Borchert et al., 2004).

The pollen-related overlap remained relatively consistent across seasons, supporting the idea that pollen constitutes a continuously available resource. Lastly, NMDS patterns for resin indicate a tendency for increased overlap during intermediate sampling periods, coinciding with peaks in resin use observed in both species. This convergence may reflect synchronized colony-level demands rather than environmental availability alone. Both species might overlap ecologically on the periods of preferred nest construction, possibly due to the unique environmental conditions of the seasonally dry tropical forests. Given that resin is a valuable and potentially limiting resource, these periods of increased overlap may also intensify competitive interactions among colonies (Shanahan & Spivak, 2021).

Overall, these findings suggest that stingless bees niche dynamics in tropical dry forests are strongly structured by seasonality, with *M. mimetica* and *Scaptotrigona* sp alternating between niche partitioning during periods of resource scarcity and convergence in resource use when resources are abundant, reflecting a dynamic balance between availability and competitive interactions that shapes coexistence in seasonal tropical systems. This is particularly interesting given that whilst in some systems niche overlap is usually tied to resource scarcity, the distinctive climatic and ecological characteristics of seasonally dry tropical forests – with periods of markedly low resources followed by periods of significantly high resource availability – have contributed to the evolution of native species in very specific and with very unique adaptations.

### Future directions

In the Tumbesian tropical dry forest and its transitional zones into Andean ecosystems, up to twelve species of stingless bees coexist, yet their trophic niches, spatial distribution, and ecological interactions remain poorly understood. A comprehensive community-level approach would allow the assessment of niche differentiation and overlap among multiple species, particularly across altitudinal gradients, providing insights into how resource use and species coexistence vary across environmental conditions. In addition, future studies should integrate ecological and socio-economic perspectives. Stingless bees are frequently managed by rural communities, often under non-sustainable practices. Investigating how native stingless bees contribute to crop pollination across different altitudinal zones and identifying the plant species that benefit most from their activity, could support the development of sustainable meliponiculture practices. Such approaches may enhance local livelihoods while promoting the conservation of native pollinator communities. Another key research priority concerns interactions with the introduced species *Apis mellifera*, particularly regarding potential competition with native species. Understanding the extent to which this introduced species may impact native pollinators is essential, especially in the context of global pollinator declines that greatly affect native species. Future work should further explore the ecological roles of stingless bees, including their foraging dynamics and contribution to ecosystem functioning. Such research would improve our understanding of biodiversity in tropical dry forests and support conservation and sustainable management strategies.

## Conclusion

This study demonstrates that niche overlap between *Melipona mimetica* and *Scaptotrigona* sp. peaks during the wet season in the seasonally dry tropical forests of Ecuador, when floral resources are most abundant. This convergence on nutritionally rich floral resources suggests an adaptive response to strong environmental seasonality and challenges the prevailing expectation that competition intensifies primarily under resource limitations.

These findings indicate that, in highly seasonal ecosystems, resource abundance may also structure competitive interactions by concentrating foraging on high-reward plant species, highlighting the role of nutritional optimization in shaping pollinator behavior. Such dynamics emphasize the importance of environmental context in refining general theories of niche partitioning and competition. Importantly, the addition of external pressures (including competition with Apis mellifera and non-sustainable management practices) may further intensify competition during critical periods of resource use. In systems where species are already finely adapted to extreme seasonal variability, these pressures may disproportionately affect native populations.

Overall, this study underscores the unique and context-dependent ecology of stingless bees in seasonally dry tropical forests and highlights their vulnerability to anthropogenic change. It emphasizes the need to incorporate seasonal resource dynamics into conservation and management strategies in these highly variable ecosystems.

## Notes

### Competing Interest Statement

The authors have declared no competing interest.

